# The short chain fatty acid propionic acid activates the Rcs stress response system partially through inhibition of D-alanine racemase

**DOI:** 10.1101/2022.08.17.504360

**Authors:** Nathaniel S. Harshaw, Mitchell D. Meyer, Nicholas A. Stella, Kara M. Lehner, Regis P. Kowalski, Robert M.Q. Shanks

## Abstract

The Enterobacterial Rcs stress response system reacts to envelope stresses through a complex two-component phosphorelay system to regulate a variety of environmental response genes such as capsular polysaccharide and flagella biosynthesis. However, beyond *Escherichia coli*, the stresses that activate Rcs are not well understood. In this study, we used a Rcs system dependent luminescent transcriptional reporter to screen a library of over 240 antimicrobial compounds for those that activated the Rcs system in *Serratia marcescens*, a *Yersiniaceae* family bacterium. Using an isogenic *rcsB* mutant to establish specificity, both new and expected activators were identified including the short chain fatty acid propionic acid found at millimolar levels in the human gut. Propionic acid did not reduce bacterial intracellular pH as hypothesized for its antibacterial mechanism. Rather than reduction of intracellular pH, data suggests that the Rcs-activating mechanism of propionic acid is, in part, due to inactivation of the enzyme alanine racemase. This enzyme is responsible for D-alanine biosynthesis, an amino-acid required for generating bacterial cell walls. These results suggest host gut short chain fatty acids can influence bacterial behavior through activation of the Rcs stress response system.

## Introduction

Regulation of capsule synthesis among Enterobacterales species is performed by the Rcs system, a highly-conserved bacterial stress response system. The Rcs pathway relies on complex phosphorelay signal transduction through the interplay of multiple inner and outer membrane components including RcsC, RcsD, RcsF, IgaA, RcsB (a response regulator), and RcsB-binding accessory proteins including RcsA (1–3). RcsF interacts with outer membrane protein OmpA to monitor outer membrane integrity (4). Together these components are responsible for transcriptional regulation of genes induced by cell wall and outer membrane stresses, the scope of which is not fully understood. Known Rcs-system activators include mutations, enzymes, peptides, and drugs that affect the outer membrane directly or through inhibition of enzymes necessary for outer membrane biogenesis (4–8). In addition, mutations, enzymes, or drugs that target peptidoglycan synthesis can activate the Rcs system (7, 9–12) as well as osmotic and oxidative stress (13, 14).

While Rcs responds to many antibiotics, activation does not confer notably reduced antibiotic susceptibility. However, as Rcs activation promotes enhanced survival at near minimum inhibitory antibiotic concentrations, activation could increase persistence of bacteria in niches in the body with near inhibitory antibiotic levels that do not achieve sufficient levels to inactivate them (12). The Rcs system is also of interest as a regulator of virulence determinants including capsule, secreted toxins, flagella, and adhesion molecules (15–19). Generally, Rcs overactive mutants exhibit reduced virulence and those with defective Rcs systems demonstrate increased virulence (20–22). Therefore, it is likely that regulated control of the Rcs system is critical during infections.

For this study, *S. marcescens* was used as a model organism. *S. marcescens* is an opportunistic pathogen of the family *Yersiniaceae* and causes contact lens associated keratitis in healthy patients and nosocomial infections, such as ventilator associated pneumonia, in the immunocompromised. *S. marcescens* is associated with microbial dysbiosis in Crohn’s disease (23) and is found in numerous environmental niches such as water, soil, plants, coral, and the gut flora of mammals and insects (24, 25). *S. marcescens* mutants with an inactivated or over-activated Rcs system demonstrate shifts in the global transcriptional landscape (19, 26). Other work has demonstrated a role for the Rcs system in regulation of *S. marcescens* virulence and virulence factors including the ShlA cytolysin (27). In *S. marcescens*, the Rcs system can be activated by mutations that affect enterobacterial common antigen synthesis and by mutation of the *S. marcescens* IgaA ortholog, GumB (28, 29). Antibiotics used for empirical treatment of ocular infections activated the *S. marcescens* Rcs system and included antibiotics to which the bacteria are susceptible (ceftazidime) and resistant (cefazolin, polymyxin B, and vancomycin). Additionally, the Rcs system regulates horizontal gene transfer and the phage defense systems of *Serratia* species (30).

In this study, a luminescence reporter for Rcs-system activation was used to screen a library of over 240 compounds to elucidate the breadth of insults the Rcs system detects. This was used in a wild type (WT) and Rcs-deficient background to establish Rcs-system specific effects. The ability of one of the most abundant gut short chain fatty acids (SCFA), propionic acid, to specifically activate the Rcs system was identified and evaluated. Propionic acid, produced by the microbiota through digestion of dietary fiber, is found at high concentrations in the human gut (up to 15-26 mM) and blood (up to 75 μM) (31–33), and so likely impacts the behavior of bacteria via the Rcs or similar stress response systems.

## Results

### Screen for chemical activators of the Rcs system

A plasmid-borne luminescence reporter construct was used to screen for molecules that activate the Rcs system (Table 1, Figure S1, Supplemental Data). The previously described plasmid includes the promoter for the SMDB11_1637 open reading frame that is predicted to code for the conserved *osmB* gene (29, 34, 35). This promoter is induced by the Rcs system in *S. marcescens* and other bacterial genera (29, 34, 35). A clinical isolate of *S. marcescens* bearing a plasmid with this reporter, pMQ747, was grown in LB medium and added to Biolog Phenotypic Microarray plates PM11-20. These 96 well plates contained over 200 compounds, many of which have antimicrobial effects. These compounds were grouped partly based on work by Dunkley, et al (36) (Supplemental Data). Each compound is replicated in 3-4 different proprietary concentrations. The bacteria were added to wells at a specified culture density and grown for four hours at which point the optical density and luminescence were measured. The relative luminescence was determined by dividing the luminescence values by the optical density. The four-hour time point was optimal based empirically on prior experiments (29). As a control, culture density was measured at the initial time point to identify the compounds that altered optical density independently of bacterial growth. To test whether the effects were specific to the Rcs system, the pMQ747 *PosmB* reporter plasmid was transformed into a Δ*rcsB* mutant lacking the Rcs response regulator, RcsB. Compounds that induced a 2-fold or higher expression in the WT compared to the Δ*rcsB* strain were considered to activate the reporter in a Rcs-dependent manner. A heat map of all of the tested compounds demonstrates clear differences between the WT and Δ*rcsB* mutant and between compound groups (Figure S1). 25 compounds were found to increase expression 2-fold or higher in the WT than the control and were significantly higher than expression in the Δ*rcsB* mutant (Figure 1). This includes known activators of the Rcs system, such as polymyxin B and vancomycin, that were previously shown to activate this reporter (29), and several compounds that have not been previously shown to activate the Rcs system including cell wall and membrane acting antibiotics. Notable among these was sodium caprylate, the sodium salt of octanoic acid. This medium chain fatty acid induced an almost 20-fold increase in luminescence (Figure 1).

**Table 1.** *S. marcescens* strains and plasmids used in this study.

| Strain /<br>plasmid | Description | Source |
| --- | --- | --- |
| K904 | Wild-type *S. marcescens* keratitis isolate | (68) |
| $\Delta rcsB$ | K904 with *rcsB* ORF replaced by *mClover* | (69) |
| pMQ414 | oriRSF1010-based plasmid with PnptII-gfp | (65) |
| pMQ747 | pMQ713 with SMDB11_1637 (*osmB*) promoter -<br>*luxCDABE* | (29) |
| pMQ748 | pMQ713 with SMDB11_2817 promoter -*luxCDABE* | (29) |
| pMQ749 | pMQ713 with *umoD* promoter -*luxCDABE* | (29) |
| pMQ802 | pMQ414 with pHLuorin2 replacing *tdtomato*, codon<br>optimized for *S. marcescens* | This study |

**Figure 1.**
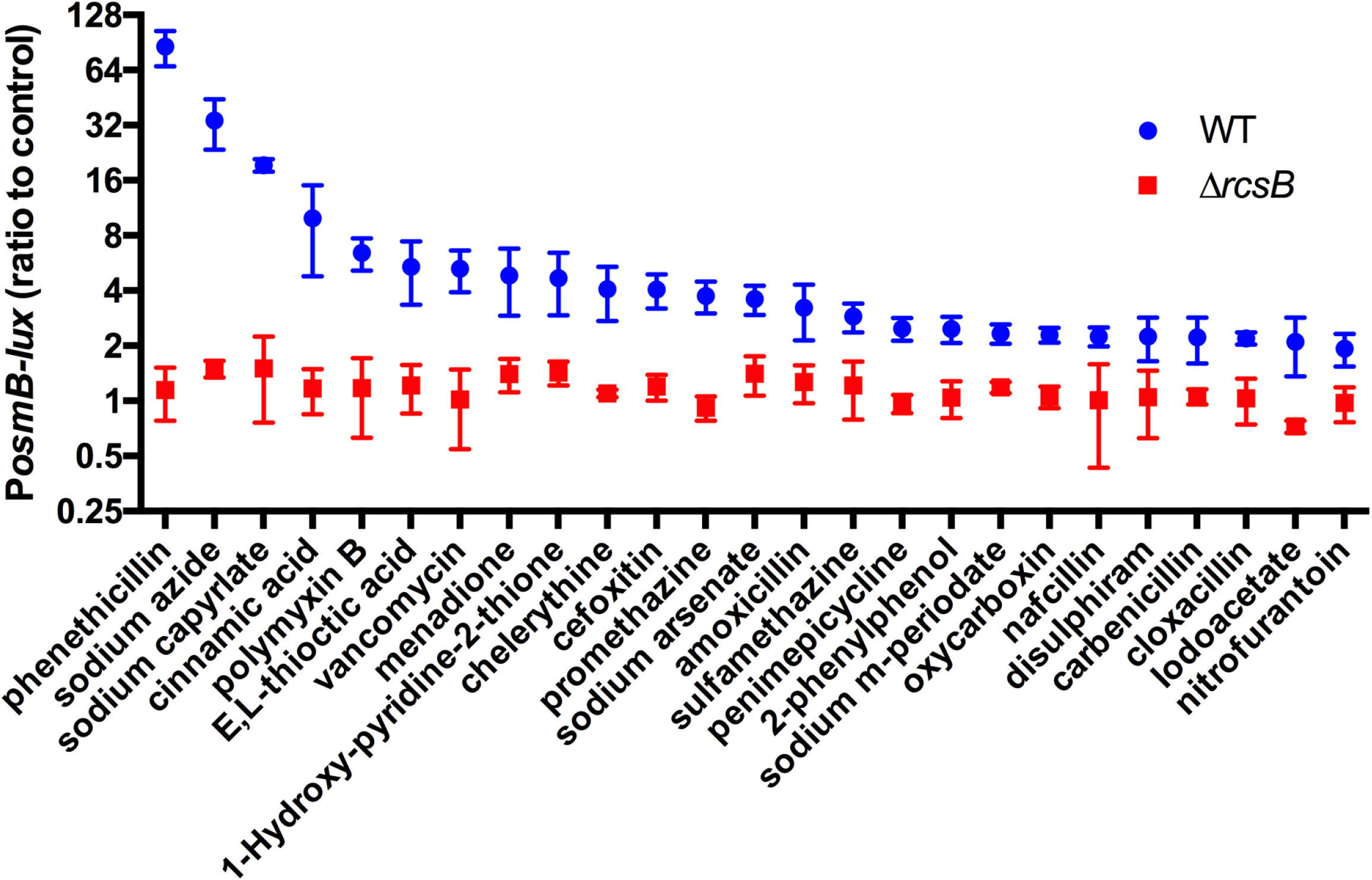
Compounds that activate an Rcs-system regulated promoter in *S. marcescens*. A luminescent reporter for the Rcs system (P*osmB*) was used to screen for compounds that activate the Rcs system from Biolog Phenotypic Microarray plates PM11-20. Only compounds that elicited a 2-fold or greater difference between WT and Δ*rcsB* are shown, p<0.05 by Student’s T-test. Mean and SD are shown, n=3.

Biolog Gen III plates were also used with the WT only. The Gen III plate consists of a 96 well plate with different potentially stress-inducing compounds, such as sodium chloride and antibiotics, designed to differentiate bacterial species. Of these compounds, three fatty acids, sodium butyrate (2.9-fold), a-keto-butyric acid (2.4-fold), and propionic acid (2.0-fold), correlated with induced reporter activity in the WT compared to the control (Figure S2).

Unexpectedly, a group of compounds activated the reporter more highly in the Δ*rcsB* mutant (Table 3, Figure S3A). We hypothesize these compounds activate other stress response systems capable of regulating the *PosmB* promoter that are repressed by the Rcs system (Figure S3B). These compounds include nucleic acid metabolism targeting antibiotics such as rifamycin SV, cinoxacin, and nalidixic acid. Two macrolide antibiotics that target translation, tylosin and spiramycin, were also identified.

### Propionate activates RcsB-regulated promoters

SCFAs were chosen for further analysis based on the data from the screen. The SCFAs were prioritized due to their high prevalence in the human gut and bloodstream, and the lack of information on their impact on the Rcs system. The *PosmB-luxCADBE* construct was used to test four different SCFAs (Figure 2). Formic acid and acetic acid had little impact on the reporter whereas butyric acid and propionic acid strongly activated expression at subinhibitory SCFA levels. The highest activation was at 6.25 mM SCFA concentration. Notably, the activation was largely, or entirely absent in a Δ*rcsB* mutant, indicating a role for the Rcs system in this process.

**Figure 2.**
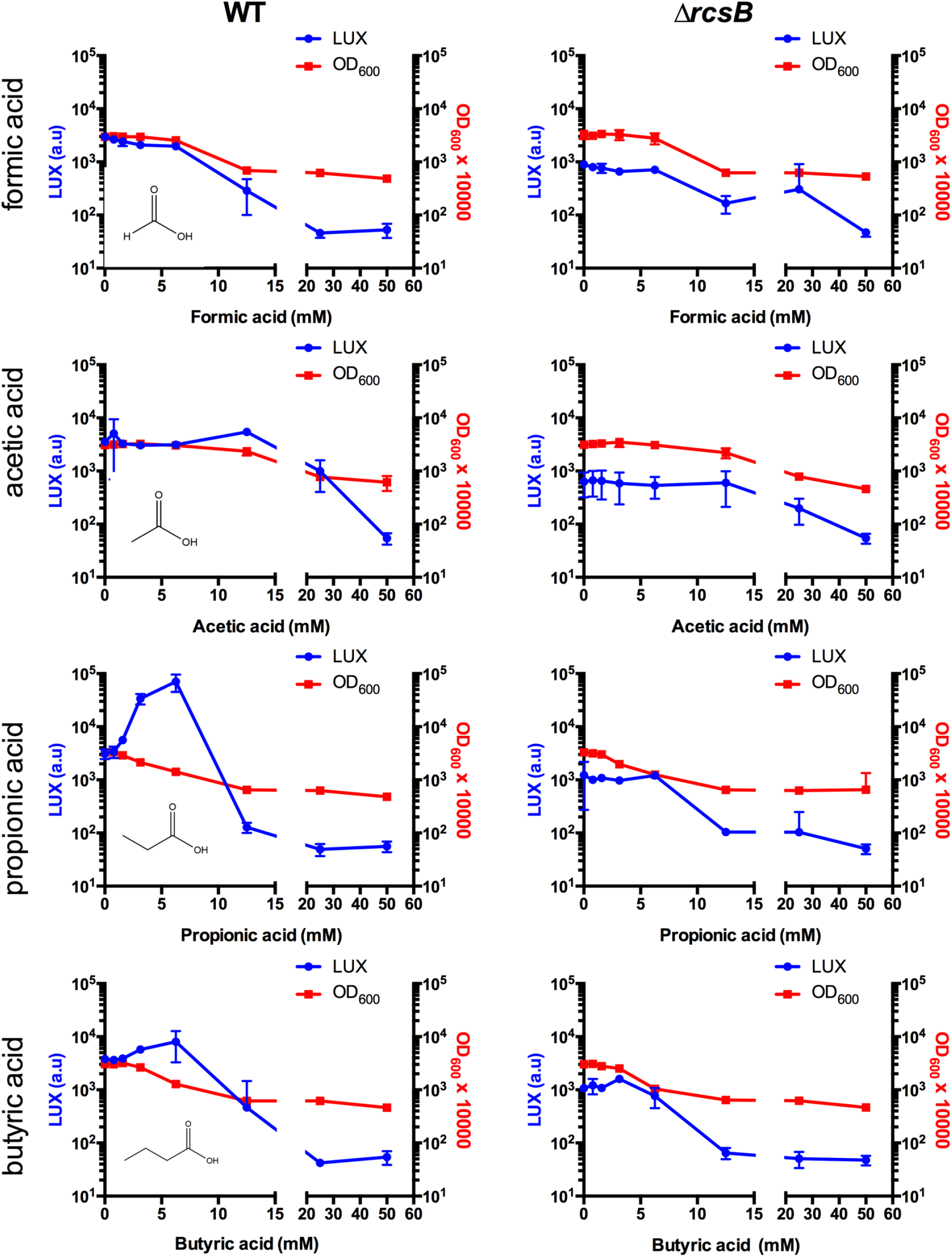
The short chain fatty acid propionate activates the Rcs system. *S. marcescens* wild type and an isogenic Δ*rcsB* mutant were evaluated for P*osmB*-driven luminescence in the presence of short chain fatty acids after co-incubation for 4 hours. Optical density at 600 nm was also determined. Propionic acid and butyric acid induced luminescence at subinhibitory concentrations in the wild type but not the Δ*rcsB* mutant. Mean and standard deviations are shown, n◻5.

To test whether this effect was common to other Rcs-regulated promoters, we selected two that were recently shown to activate Rcs in this strain (29). The promoters, P_*SMDB11_2817*_ and P*umoB*, were also highly responsive to propionic acid in the WT background, but minimally or not activated in the Δ*rcsB* mutant (Figure 3).

**Figure 3.**
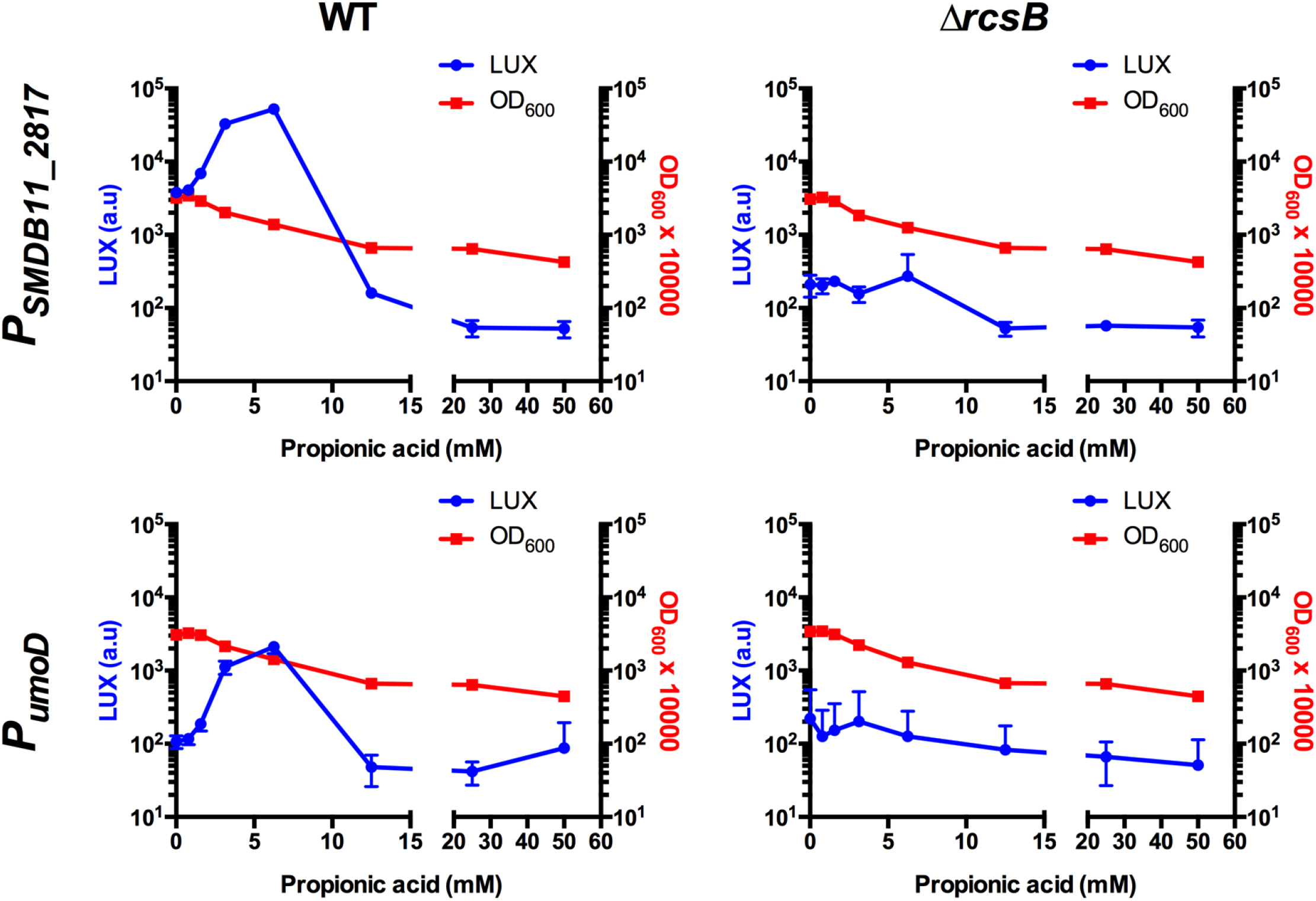
Propionic acid activates Rcs-dependent promoters P_*SMDB11_2817*_ and P_*umoD*_. *S. marcescens* strains with two different RcsB-influenced promoters driving *luxCDABE* produced increased luminescence in the presence of propionic acid after co-incubation for 4 hours. Optical density at 600 nm was also determined. Means and standard deviations are shown, n≥6. This indicates that the Rcs-dependent effect of propionic acid is common to multiple Rcs activated promoters.

### Propionic acid mechanism of growth inhibition and Rcs activation

The previously hypothesized mechanism by which propionic acid inhibits bacterial growth has been through reduction in intracellular pH (37). Undissociated forms of organic acids readily penetrate into bacterial cells, where they dissociate, thus impeding pH homeostasis (38). To test whether propionic acid concentrations that induce the Rcs system in *S. marcescens* caused a reduction in intracellular pH, we used a pH sensitive GFP variant, pHLuorin2 (39). The construct was validated as producing fluorescence in a pH dependent manner (R^2^=0.943) and was responsive to culture acidification through growth with glucose (2% w/v). This reduced the intracellular pH from 7.1 in LB medium to 6.5 in LB with glucose (Figure S4) which is below 6.8, the reported intracellular pH which inhibits bacterial growth (40).

Increasing the propionic acid in the medium prior to inoculation resulted in a corresponding drop in pH of the LB media, decreasing from 6.9 with no propionic acid to 5.7 with 6.25 mM (Figure 4). In the absence of propionic acid, the growth medium pH increased above pH 8 as expected for *S. marcescens* (41, 42). In tubes with a propionic acid concentration of 6.25 mM the pH lowered to 7.7 (Figure 4). The intracellular pH was calculated to be between 7.0 and 7.1 regardless of propionic acid concentration.

**Figure 4.**
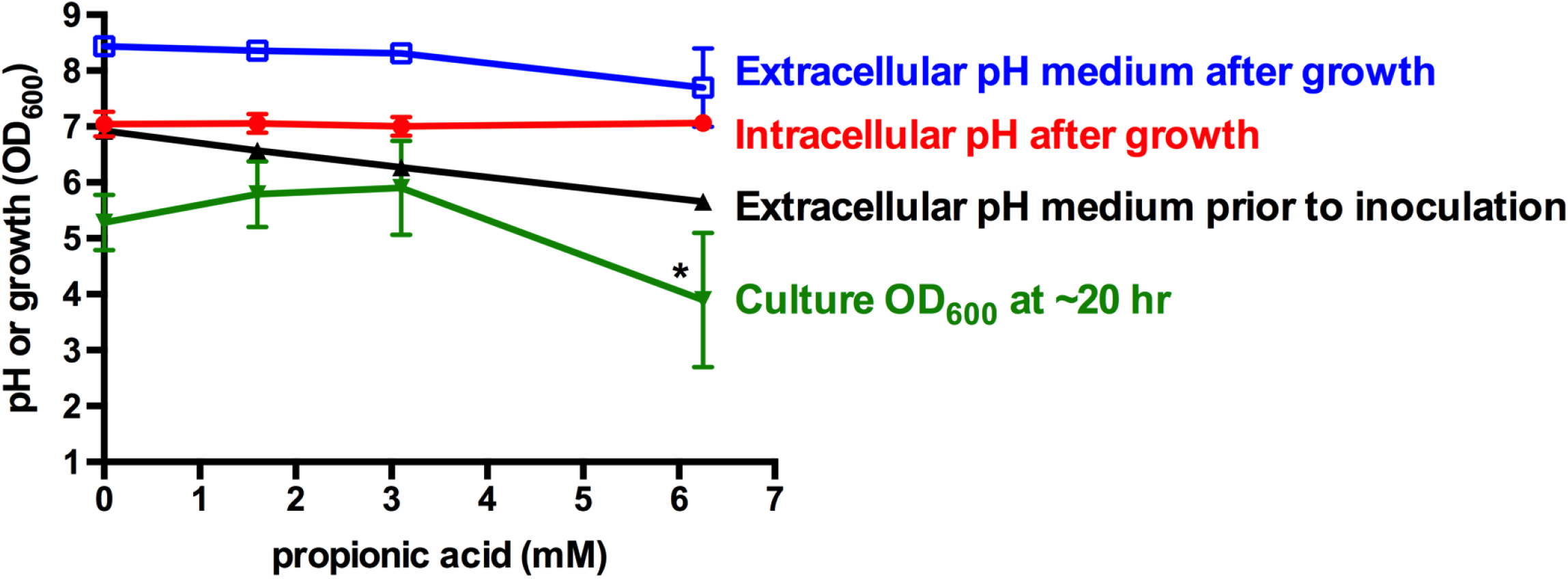
Propionate does not activate the Rcs system through lowering intracellular pH. A pH sensitive GFP variant, pHIuorin2 was codon optimized for *Serratia marcescens* and used to assess intracellular pH. Fluorescence was measured after overnight growth in LB broth in the presence and absence of propionate. Propionic acid at concentrations that activate the Rcs system did not alter intracellular pH (red circles). Medium pH prior to bacterial inoculation was reduced with increasing propionic acid concentration (black triangles). Medium pH after growth was made alkaline by bacterial growth effects (blue squares). Culture density was reduced when bacteria were grown with 6.25 mM propionic acid, but not lower concentrations (green inverted triangles). Asterisk indicates significantly different from 0 mM propionic acid (ANOVA, Tukey’s post-test, p<0.05). n=4, mean and SD are shown.

### Activation of Rcs through inhibition of alanine racemase

These data suggest that it is unlikely that propionic acid activates the Rcs system through acidification of bacterial cells’ intracellular environment. Propionate is a known inhibitor of alanine racemase for *Bacillus stearothermophilus* (43, 44), *Staphylococcus aureus* (45), and *Streptomyces coelicolor* (45). Alanine racemase is an essential enzyme for bacterial cell wall biosynthesis through conversion of L-alanine to D-alanine, which is incorporated into the peptidoglycan cell wall (43, 46). Inactivation of alanine racemase prevents cell wall biosynthesis, which would be expected to activate the Rcs system and lead to bacterial cell death. If this is true, then other alanine racemase inhibitors should activate the Rcs system. D-cycloserine is an inhibitor of both alanine racemase and D-alanine ligase. The P*osmB*-*lux* reporter was activated (>30%) in the WT by D-cycloserine at the maximum tolerated concentration of 25 mg/ml (245 μM) compared to no D-cycloserine (Figure 5A). By contrast, the reporter was not activated in the Δ*rcsB* mutant (Figure 5A). Another D-alanine racemase inhibitor, β-chloro-D-alanine was used, and induced a stronger increase (69%) in expression from the P*osmB*-*luxCDABE* reporter at 25μg/ml (156 μM) in the WT but not in the Δ*rcsB* mutant (Figure 5B).

**Figure 5.**
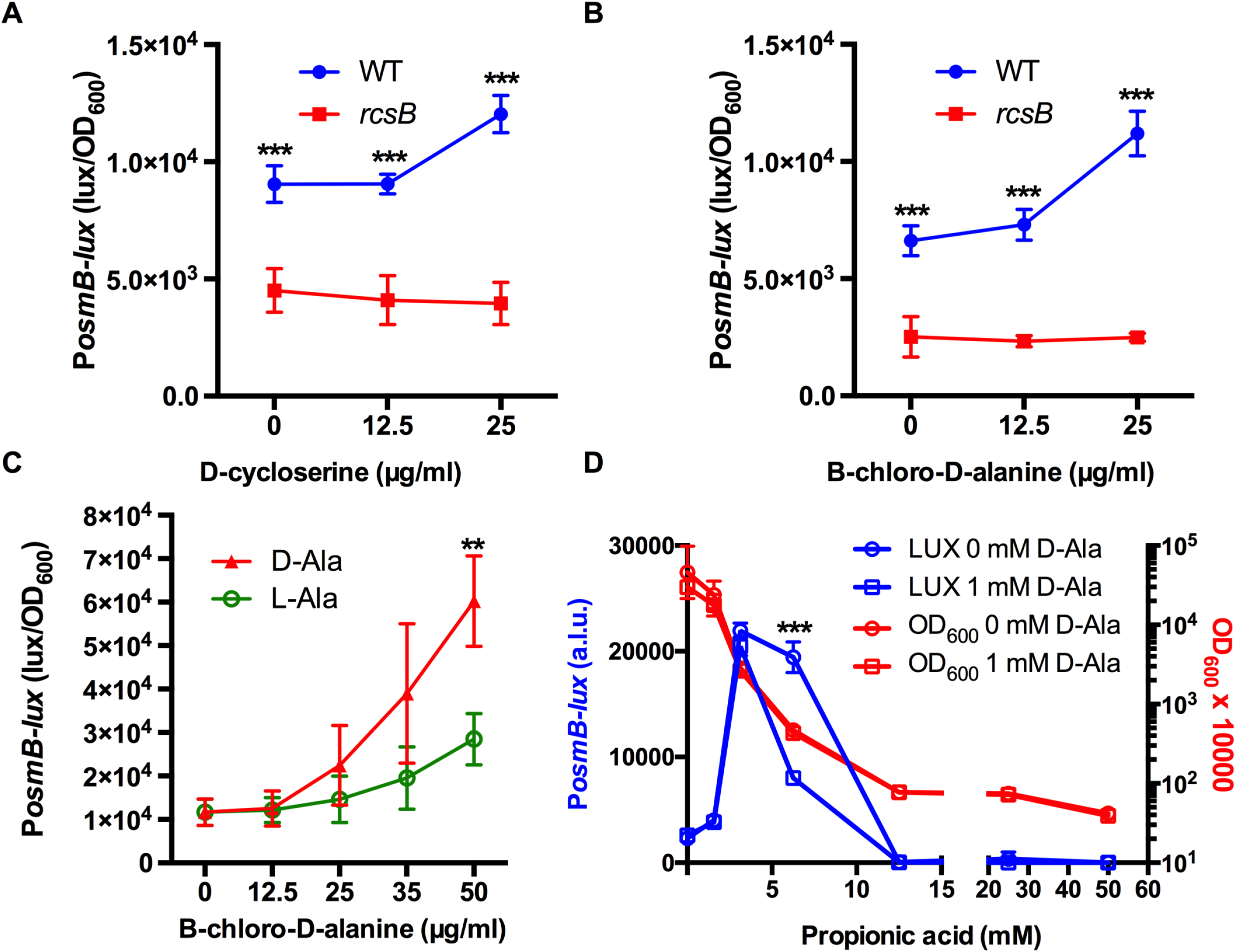
Alanine racemase inhibitors activate the Rcs system. Bacteria were grown in LB media with D-alanine racemase inhibitors for 4 hours at 30°C. P*osmB*-driven luminescence values normalized by culture density at 4 hours post inoculation. **A**. Luminescence response to D-cycloserine. **B**. Luminescence response to β-chloro-D-alanine hydrochloride. **C**. D-alanine but not L-alanine (1 mM) reduced the induction of *osmB* by β-chloro-D-alanine in the WT. **D**. D-alanine reduced the luminescence activation by propionic acid. Mean and SD are shown, n≥4. **, p<0.001; ***, p<0.0001.

If the Rcs-dependent increase in *PosmB*-*lux* reporter by propionic acid is triggered by inhibition of the D-alanine racemase enzyme, then exogenous D-alanine would be expected to reduce cellular stress corresponding with lower luminescence values. To test this, bacteria were subject to a range of Beta-chloro-D-alanine from 0 to 50 μg/ml in both the presence and absence of D-alanine or L-alanine (1 mM) (Figure 5C). Similarly, D-alanine reduced *PosmB* reporter activity of 6.25 mM propionic acid challenge (Figure 5D). However, consistent with propionic acid affecting cells in multiple ways, D-alanine did not rescue growth inhibition by propionic acid. Together, these data suggest that propionic acid activates the Rcs system partially through inhibition of the D-alanine racemase enzyme, and that inhibition of growth likely involves multiple targets for propionic acid and is not solely due to inactivation of D-alanine racemase caused by intracellular acidification.

### Inhibition of *S. marcescens* alanine racemase activity by propionic acid *in vitro*

A prerequisite for our model is that propionic acid can inhibit the *S. marcescens* alanine racemase. While inhibition of alanine racemase has been demonstrated for certain Gram-positive bacteria, it has not been demonstrated with Gram-negative bacteria such as *S. marcescens*. An enzymatic approach was used to ascertain alanine racemase activity in partially purified wild-type lysates from which compounds smaller than 10 kD, such as D-alanine, were removed (see materials and methods and Figure 6A). The experiments indicated that the *S. marcescens* lysates contained robust alanine racemase activity (Figure 6B).

**Figure 6.**
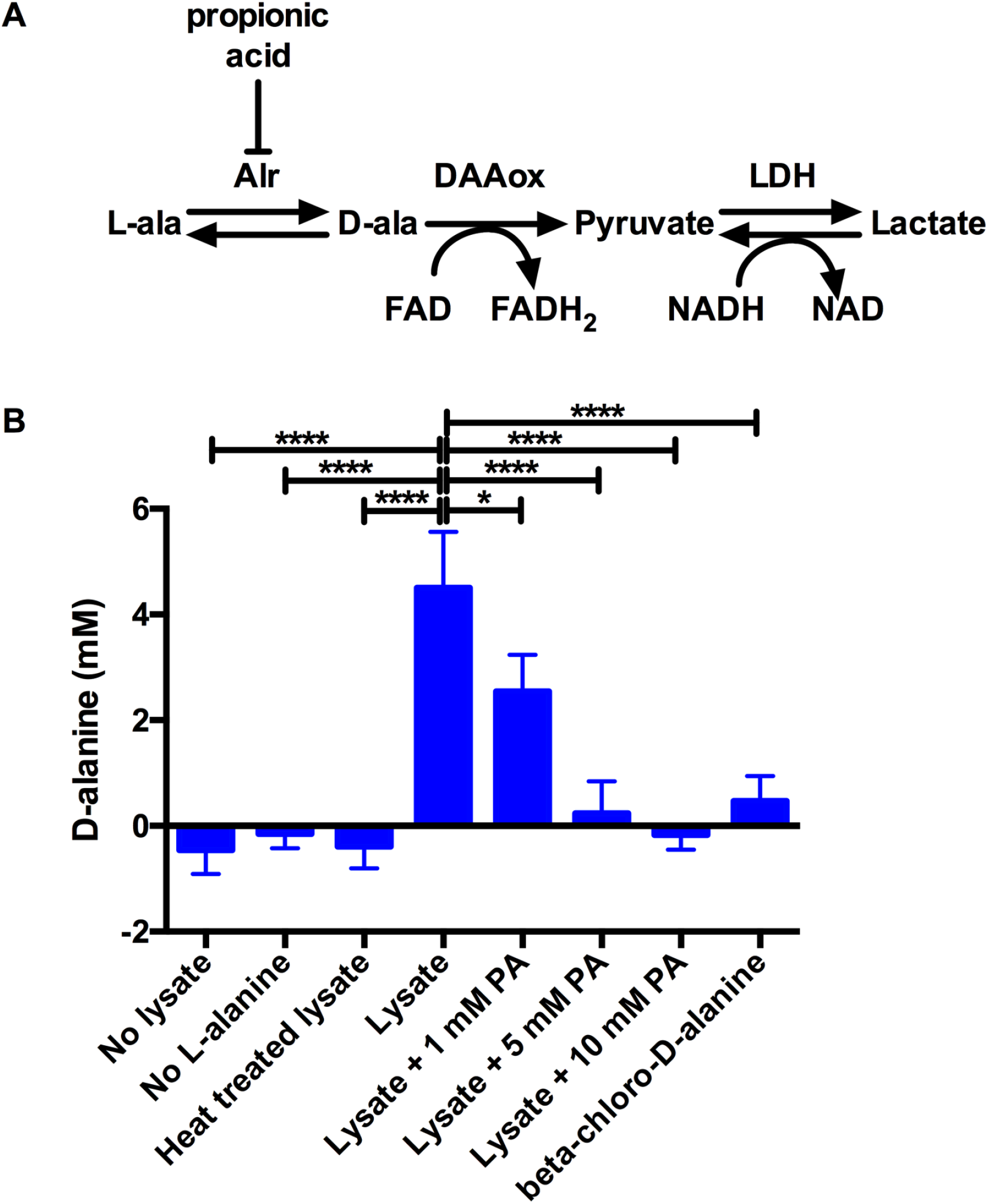
Propionic acid (PA) inhibits *S. marcescens* alanine racemase activity in vitro. **A**. Schematic for the enzymatic activity for conversion of L-alanine (L-ala) used in this assay. The alanine racemase activity (Alr) was provided from WT *S. marcescens* lysates. The reaction was measured by the change in NADH levels as measured by absorbance at 340 nm. DAAox is D-amino acid oxidase; LDH is lactate dehyrogenase. **B**. D-alanine concentration, serves as an indicator for alanine racemase activity and this activity as present in the lysate, but reduced by propionic acid or alanine racemase inhibitor beta-chloro-D-alanine. L-alanine at 10 mM was introduced into the experiment and its conversion to D-ala was determined by comparison with a standard curve. Significance was determined using ANOVA with Tukey’s post test. *, p<0.05, ****, p<0.0001. Mean and SD are shown, n=3.

To evaluate whether propionic acid inhibited the *S. marcescens* alanine racemase activity *in vitro*, propionic acid was added to the lysate and was found to inhibit alanine racemase in a dose dependent manner (Figure 6B). Similarly, alanine racemase inhibitor β-chloro-D-alanine inhibited the reaction as expected (Figure 6B). As a control, it was determined that propionic acid did not inhibit other enzymes required for the analysis, indicating that the inhibitory effect was on alanine racemase (Figure S5). Together these data suggest that *S. marcescens* alanine racemase activity can be inhibited by propionic acid and support the model that propionic acid activates the Rcs system partly through inhibition of alanine racemase activity.

### Propionate impacts flagella-based motility and *fliC* gene expression

The bacterial flagellum is a potent activator of inflammation through binding with TLR5 (47). Flagellar genes are regulated by the Rcs system in numerous species including *S. marcescens* (19, 26). We tested whether physiologic concentrations of propionic acid had an impact on *S. marcescens* flagellum-based motility (Figure 7). An mClover transcriptional reporter for the *fliC* promoter increased over time in the WT, so an overnight (16h) culture was used to evaluate the effect of propionate. For the WT, increasing propionate levels correlated with reduced fluorescence even when growth was largely unaffected (Figure 7A). The Δ*rcsB* mutant was not used because it has the *rcsB* gene replaced by *mClover* and we could not distinguish between P*fliC* and P*rcsB* mediated mClover fluorescence. Swimming motility was evaluated to determine whether this change in *fliC* expression in the WT conferred a phenotype (Figure 7B). Both the WT and Δ*rcsB* mutant were inhibited for swimming zones, with a greater effect on the WT (Figure 7B). As expected, swarming motility was distinctly greater in the Δ*rcsB* mutant due to derepressed flagellar operons (48–50), and both genotypes had swarming partially inhibited by propionic acid at 6.25 mM (Figure 7C). These data suggest that propionic acid can inhibit flagellar based motility both by Rcs-dependent and independent mechanisms.

**Figure 7.**
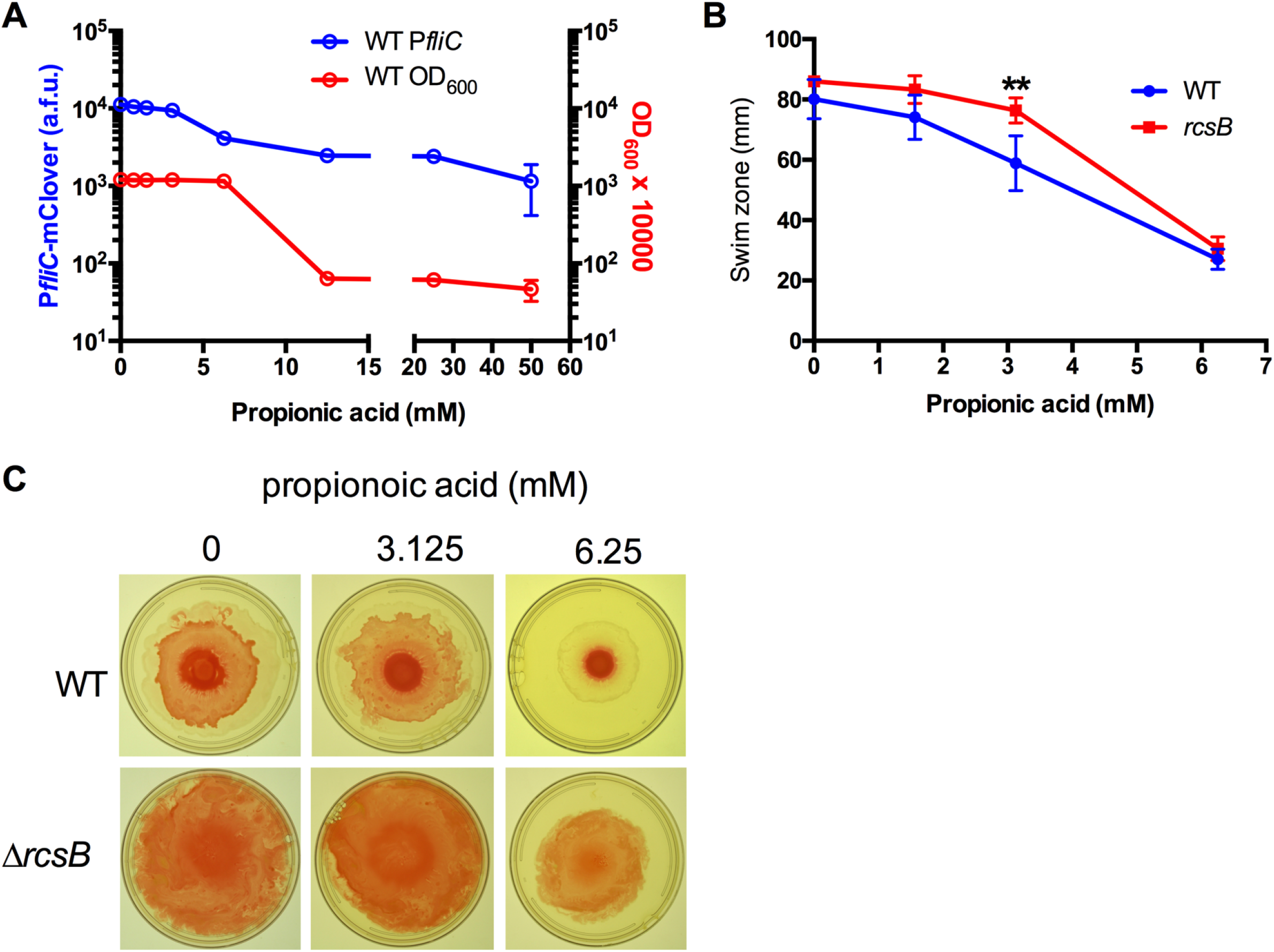
Regulation of *fliC* and flagella-based motility by propionic acid. **A**. Fluorescence from *fliC* transcriptional reporter and corresponding optical density, after 16 h at 30°C, n=16. **B**. Swim zone at 16h at 30°C, LB with agar at 0.3% (w/v). **, p<0.01. n≥5. **C**. Representative images of swarming motility on LB agar (1% agar).

## Discussion

### Rcs system is activated by propionic acid

Our screen for compounds that induce the Rcs system has identified new activators, notably the short chain fatty acid propionic acid. In some cases, Rcs-activated promoters can be controlled by other stress response systems (7). Strong support for propionic acid activating the Rcs system is its ability to activate three different Rcs-regulated promoters in the WT, but not an isogenic Rcs-deficient mutant. Due to its antimicrobial effects, propionic acid is commonly used as a food preservative (38), used topically to treat skin infections (51), and may affect antibiotic susceptibility (52, 53). However, the mechanism of action is not fully understood and is hypothesized to be derived from intracellular acidification (38). Intracellular acidification is not expected to strongly activate the Rcs system, and indeed, we saw that propionic acid did not cause intracellular acidification at concentrations that activated the Rcs system reporter. Due to these data, we tested an alternative hypothesis: propionic acid inhibits D-alanine racemase activity as was shown for several Gram-positive bacterial species. Inhibition of this key enzyme in cell wall biosynthesis would be expected to activate the Rcs system. Our data suggest that subinhibitory concentrations of known D-alanine racemase inhibitors increase expression from an Rcs dependent promoter in an RcsB-dependent manner and that exogenous D-alanine can quell this induction. Furthermore, our data support propionic acid inhibition of *S. marcescens* D-alanine racemase *in vitro*. Together, these data support a role for D-alanine racemase targeting by propionic acid contributing to activation of the Rcs system.

However, promoter activation by D-alanine inhibitors did not achieve that of propionic acid, suggesting propionic acid has additional effects on the cell membrane that the Rcs system detects. For instance, bacterial outer membrane lipid content and protein profiles can be influenced by short chain fatty acids like propionate in *Borrelia burgdorferi*, *Prevotella bryantii*, and *Pseudomonas aeruginosa* (53–55). Freese, et al. hypothesized that the antimicrobial target of short chain fatty acids is the membrane (56). These membrane effects may be responsible for propionic acid induction of the Rcs system.

SCFAs such as butryrate and propionate impact the immune system and can influence host responses to herpes simplex virus (57), the outcomes of dry eye diseases (58). Additionally, they can impact host responses through binding to the FFAR2 G-protein-coupled receptor and influencing the response to LPS (59, 60). Bacterial behavior of Gram-negative and Gram-positive is also highly influenced by SCFA (61, 62). Our study showed a measurable impact of propionic acid on flagella-based motility. As the flagellum is a major pathogen associated molecular pattern, this inhibition could influence host-pathogen interactions.

Our study also confirmed several established Rcs activating compounds and compound classes such as β-lactam antibiotics. General membrane targeting antimicrobial compounds, such as polymyxin B, were also identified. Here, we find three previously unreported activators of the Rcs system: chelerythrine, cinnamic acid, and 4-aminopyridine, molecules predicted to be transporter inhibitors, permeability change, and proton motive force uncouplers, respectively. Other predicted PMF uncouplers, ethanol and antimicrobial peptide Gramicidin A, are known to activate the Rcs system (6). We also observed that PMF uncoupler CCCP increased reporter activity ~8 fold above that of the Δ*rcsB* mutant, but this did not reach our threshold of significance (p=0.12). SCFAs may also affect membranes and membrane protein function through disruption of the proton gradient (38).

Questions remain regarding how physical damage may be responsible for activation of the Rcs system. While outer membrane perturbations can activate the system, this is not always the case. Systematic analysis by Steenhuis and colleagues demonstrated that compounds which increased outer membrane permeability (edthylenediaminetetraacetic acid (EDTA) and sodium dodecyl sulfate (SDS)) or compromised outer membrane integrity (triclosan) failed to activate the Rcs system of *E. coli* (7). The authors concluded that the Rcs system responds to the interaction of antimicrobial peptides with LPS, rather than permeabilization of the outer membrane, likely due to RcsF monitoring of the LPS layer (7). Nevertheless, the extent of Rcs activation by peptides was variable even at 0.5 x minimum inhibitory concentration values, and the difference could not be explained by the molecular weight of the tested compounds (7).

The impact of cell wall acting antibiotics on induction of the Rcs system is also similarly nuanced. A number of β-lactams in this study strongly activated the Rcs system, while others did not. This may not just be an issue of susceptibility, as the *S. marcescens* Rcs system can be activated by cell wall targeting antibiotics such as cefazolin and vancomycin for which it is resistant (29). In a study by Sailer, et al., numerous peptidoglycan antibiotics were tested for activation of the *cps* genes, a classic output for activation of the Rcs system in *E. coli*. Interestingly, despite inhibiting bacterial growth, approximately half of the tested β-lactams were unable to activate expression of a *cps* transcription (9). This was not obviously due to inhibition of specific penicillin binding proteins (PBPs), although it is clear that simultaneous mutation of several PBPs is sufficient to activate the Rcs system (63). Other differences between studies have arisen. For example, ampicillin causes induction of a fluorescent reporter for the Rcs regulated *rprA* gene (7) and increased expression of Rcs controlled genes as determined by microarray in *E. coli* (10). By contrast, in our study, ampicillin induced a modest 30% induction of an Rcs-dependent promoter in the WT compared to the Δ*rcsB* mutant. Some differences in study outcomes may be due to the difference in the sensitivity of the assay, time point of the analysis, concentration of drug tested, and differences between species with respect to drug permeability. Alternatively, the different effects may indicate that highly specific, rather than general, insults are detected by the Rcs system.

In conclusion, this study identified the SCFA propionic acid as a strong inducer of the Rcs stress response system of *S. marcescens*. Due to the important role of the Rcs system in controlling bacterial behaviors such as virulence factor production, this suggests a role for SCFAs in the gut in programming enteric bacteria to reduce virulence factor production. The Rcs system is, in general, highly conserved among the tested Enterobacterales, with notable exceptions such as *Yersinia pestis*, which has a highly active RcsC, highlighting the need for validation in other gram negative species (64). This study also underscores the potential of propionic acid as an alternative to antibiotics that also reduce virulence factor production.

## Materials and Methods

### Reporter Assays

Glycerol stocks of *S. marcescens* strains were maintained at −80°C and are listed in Table 1. All incubation of *S. marcescens* were performed at 30°C in lysogeny broth (LB) with or without agar. The LB medium was supplemented with 10 μg/mL gentamicin to maintain only cells that contain the integrated lux plasmid. To obtain single colonies, *S. marcescens* strains were grown up overnight on LB agar streaking for isolation. Single colonies were then grown up overnight in LB with gentamicin, receiving aeration on a tissue culture roller. After ~18h, overnight cultures were normalized in LB with gentamicin to OD_600_ = 0.1. 100 μL of normalized culture were added to each well of Biolog GenIII and Phenotype MicroArray 11-20 MicroPlates™. The plates were rocked for 5 minutes to assure proper dissolution of plate compounds. Plates were placed in a plastic bag with a wet paper towel and incubated at 30°C. Wells were transferred to a black-sided, clear-bottomed 96 well plates (Nunc 165305) to prevent luminescence cross-contamination from one well to the other. Lux and OD_600_ readings were taken after four-hour, and in some cases six-hour, incubation time. Relative Lux values were calculated by dividing the lux value by the OD_600_. These values were then normalized to the control well to determine the fold change in Lux expression caused by each plate compound. In some cases, the ratio of expression of reporter activity in the WT to Δ*rcsB* was also measured.

*mClover* reporter assays were done as above except that fluorescence was measured from bacteria in black-sided, clear-bottomed 96 well plates (Nunc 165305) with a plate reader (Biotek Synergy 2) using 485/20 nm excitation and 516/20 nm emission filters. Optical density at 600 nm was also measured. Fluorescence was measured after overnight growth (~18 hr).

### Cloning and intracellular pH analysis

The pH sensitive *gfp* variant, pHluorin2 (39), was codon optimized for *S. marcescens* using online software (Integrated DNA Technologies) and placed under control of the constitutive *nptII* promoter. A synthetic DNA fragment with the pHluorin2 open reading frame was designed to recombine with fluorescent protein expression vector pMQ414 (65) using yeast in vivo cloning (66). The sequence from 5’ to 3’ is ggcgtttcacttctgagttcggcatggggtcaggtgggaccaccgcgctactgccgccaggcaaattctgttttatcagaccgctt ctgcgttctgatttaatctgtatcaggatccTTTATACAGTTCGTCCATGCCGTGGGTTATGCCCG CAGCAGTCACAAACTCGAGCAGCACCATGTGGTCCCGCTTTTCATTGGGGTCT TTTGAGAGGGCCGATTGGGTGTGGAGGTAGTGGTTGTCTGGCAACAGTACTG GACCATCCCCAATTGGAGTGTTTTGCTGATAATGGTCGGCAAGCTGTACGCTG CCATCTTCAATATTATGGTGCACTTGAAAAATGGCCTTTGTTCCATTTTTCTGC TTGTCCGCCATGATATACACGAGATGTTCATTATAGTTGTACTCGAGTTTATG CCCCAGTATGTTACCGTCTTCCTTAAAGTCTATCCCTTTGAGTTCTATCCGATTTACCAACGTGTCCCCCTCGAACTTTACCTCGGCGCGTGTCTTGTAGTTCCCGT CATCCTTGAAGAATATAGTCCGCTCCTGGACATATCCTTCGGGCATTGCAGAC TTGAAGAAGTCATGCTGCTTCATATGATCGGGGTAGCGGGAAAAACACTGGA CTCCGTACGACAGTGTTGTGACGAGCGTGGGCCACGGAACTGGCAATTTTCC GGTCGTGCATATAAATTTGAGGGTCAACTTACCATATGTTGCGTCACCCTCTC CTTCTCCCGAAACCGAAAATTTGTGCCCGTTAACATCACCGTCCAATTCGACG AGTATAGGGACCACACCTGTGAACAATTCCTCGCCTTTGCTCATgaattctcctcatcctgtctcttgatcagatcttgatcccctgcgccatcagatccttggcggcaagaaagccatccagtttactttgcagggcttcccaacc ttaccagagg. The pMQ414 vector was cut with BamH1 and EcoR1 to remove tdtomato and moved into *Saccharomyces cerevisiae* strain InvSc1 with the synthetic DNA as previously described (66). The resulting plasmid, pMQ802, was verified by PCR and junctions were sequenced. The plasmid was moved into *S. marcescens* strain K904 by conjugation.

Intracellular pH was determined as previously described (39). First, a standard curve was established using buffered medium. Cultures of *S. marcescens* with plasmid pMQ802 (see below) were grown in 5 mL LB for 16 hours at 30°C with aeration and subcultured 1:100. These were grown until they achieved OD_600_= 2. 1 mL aliquots of normalized subcultures (OD_600_= 2) were decanted into microfuge tubes and pelleted by centrifugation. Pellets were resuspended in 1 mL of buffers (50 mM) at pH 5.5 MES (Sigma, product M-5057), 7.0 MOPS (Sigma, product M1254), and 8.3 TAPS (Sigma, product T5130), along with 50 mM methylamine HCl (Sigma, product M0505) and 50 mM sodium benzoate (ThermoFischer, Heysham, Lancashire, LA3 2XY, UK, product A15946). 0.2 mL aliquots were transferred into wells of a black-sided, clear-bottomed 96-well plate (ThermoFisher, Waltham, MA, USA, product 165305). The OD_600_ was recorded as well as fluorescence with a plate reader (Molecular Devices SpectraMax M3, San Jose, CA, USA). Fluorescence measurements were at excitation/emission wavelengths of 485/515 and 405/515, and then the ratio of fluorescence at excitation 405 over 485 (R_405/485_) was calculated for construction of a calibration curve. These were compared to a buffer blank and a linear standard curve was established.

To test the effect of propionic acid (various concentrations) and glucose (2% w/v) on the fluorescent activity of the pH dependent GFP, cultures of *S. marcescens* with plasmid pMQ802 were started from −80°C glycerol stocks with 5 mL LB, 5 μL gentamicin, and a specified concentration of PA or glucose. The concentrations of PA supplemented growth media were 1.6 mM, 3.1 mM, and 6.3 mM (Sigma, product P1386). The effect of PA was measured with the same procedure used in the verification of pH dependence of *S. marcescens* with plasmid pMQ802. However, during resuspension, the pellets were resuspended in 1 mL of a solution containing 50 mM methylamine HCl and 50 mM sodium benzoate. 200 μL aliquots from the resuspended cultures were organized into wells of a black 96-well clear-bottomed plate. The OD_600_ was measured as well as fluorescence at 485/515 and 405/515 and ratios were compared to the standard curve.

### Literature search

To identify whether compounds have been previously determined as activating the Rcs system, the following searches were taken: 1) Google Scholar “Rcs system” and “compound name”, 2) Google Scholar “RcsB” and “compound name”, 3) NCBI PubMed “Rcs system” and “compound name”, 4) NCBI PubMed “RcsB” and “compound name”. Compounds were listed as not previously reported if no specific description of the compound activating transcription of Rcs-dependent genes was previously described based on the literature search. These are listed in Table 2.

**Table 2.** Compounds that induce the *PosmB-luxCDABE* reporter.

| Compound | Plate | Compound<br>class | Function | Fold WT<br>compound /<br>control | Fold<br>(WT/$\Delta$<br>*rscB*) | p-value |
| --- | --- | --- | --- | --- | --- | --- |
| phenethicillin | PM19 | Beta-lactam | Cell Wall<br>Targeting | 86.0 | 74.9 | 0.001 |
| sodium azide | PM15 | Inhibitor of<br>oxidative<br>phosphorylation | Metabolic<br>Compounds/Effe<br>ctors | 34.0 | 22.7 | 0.006 |
| sodium<br>caprylate | PM19 | Medium chain<br>fatty acid | Other | 19.4 | 12.9 | 0.000 |
| cinnamic acid | PM19 | Cell membrane<br>effector | Transport<br>Inhibitors/permea<br>bility<br>change/PMF<br>Uncoupling | 9.9 | 8.5 | 0.042 |
| polymyxin B | PM19 | polymyxin | Cell Wall<br>Targeting | 6.4 | 5.5 | 0.003 |
| E,L-thioctic<br>acid | PM19 | Redox<br>compound | Redox<br>Compound | 5.3 | 5.2 | 0.007 |
| vancomycin | PM12 | Glycopeptide<br>antibiotic | Cell Wall<br>Targeting | 5.4 | 4.4 | 0.025 |
| menadione | PM15 | medication | Repurposed<br>Medication | 3.7 | 4.1 | 0.003 |
| 1-hydroxy-<br>pyridine-2-thione | PM14 | chelator | Chelating Agents and Surfactants | 4.1 | 3.7 | 0.018 |
| chelerythine | PM14 | Permeability effector | Transport Inhibitors/permeability change/PMF Uncoupling | 4.8 | 3.5 | 0.038 |
| cefoxitin | PM14 | Cephalosporin | Cell Wall Targeting | 4.0 | 3.4 | 0.005 |
| promethazine | PM14 | Medication/ Metabolic inhibitor | Repurposed Medication | 4.7 | 3.3 | 0.033 |
| sodium arsenate | PM14 | General toxin | Other | 2.1 | 2.9 | 0.032 |
| amoxicillin | PM11 | Beta-lactam | Cell Wall Targeting | 2.5 | 2.6 | 0.002 |
| sulfamethazine | PM12 | Sulfonamide | Nucleic Acid Synthesis | 3.6 | 2.6 | 0.007 |
| penimepicycline | PM12 | tetracycline | Protein Synthesis | 3.2 | 2.5 | 0.039 |
| 2-phenylphenol | PM18 | disinfectant | Disinfectants | 2.9 | 2.4 | 0.012 |
| sodium m-periodate | PM18 | Redox compound | Redox Compound | 2.5 | 2.4 | 0.006 |
| oxycarboxin | PM17 | Inhibitor of oxidative phosphorylation | Metabolic Compounds/Effectors | 2.2 | 2.2 | 0.027 |
| nafcillin | PM11 | Beta-lactam | Cell Wall Targeting | 2.3 | 2.2 | 0.001 |
| disulphiram | PM19 | medication | Repurposed<br>Medication | 2.2 | 2.2 | 0.047 |
| carbenicillin | PM12 | Beta-lactam | Cell Wall<br>Targeting | 2.2 | 2.1 | 0.004 |
| cloxacillin | PM11 | Beta-lactam | Cell Wall<br>Targeting | 2.2 | 2.1 | 0.033 |
| iodoacetate | PM14 | Protein synthesis<br>inhibitor | Protein Synthesis | 1.9 | 2.0 | 0.020 |
| nitrofurantoin | PM14 | Nucleic acid<br>synthesis<br>inhibitor | Nucleic Acid<br>Synthesis | 2.3 | 2.0 | 0.002 |

a. Biolog Plate number
b. Average of WT levels with compound over WT levels in control wells. N=3
c. Average of WT levels divided by $\Delta rcsB$ mutant levels. N=3
d. Student's T-test of WT versus $\Delta rcsB$ levels.

**Table 3.** Compounds that induce the *PosmB-luxCDABE* reporter more highly in the Δ*rcsB* mutant.

| Compound | Compound class | Function | Fold $\Delta rcsB$ compound / control | Fold ($\Delta rcsB$/ WT) | p-value |
| --- | --- | --- | --- | --- | --- |
| rifamycin SV | Rifamycins | Nucleic Acid Synthesis Inhibitor | 35.4 | 8.4 | 0.001 |
| Boric acid | Antiseptic, disinfectant | Antiseptic, Disinfectants | 23.8 | 13.5 | 0.033 |
| Sodium metaborate | Metal Ions | Possible oxidative metabolism inhibitor | 20.9 | 8.6 | 0.003 |
| cinoxacin | Quinolone | Nucleic Acid Synthesis Inhibitor | 16.4 | 6.7 | 0.002 |
| 3,5-diamino-1,2,4-triazole (Guanazole) | Cytostatic triazole | Nucleic Acid Synthesis Inhibitor | 12.0 | 6.1 | 0.037 |
| Nalidixic Acid | Quinolone | Nucleic Acid Synthesis Inhibitor | 8.6 | 2.5 | 0.045 |
| DL-methionine hydroxamate | Amino acid analog | Metabolic Compound/Inhibitor | 5.3 | 3.0 | 0.012 |
| Tylosin | Macrolide | Protein synthesis inhibitor | 4.4 | 3.6 | 0.026 |
| Oxolinic acid | Quinolone | Nucleic Acid<br>Synthesis<br>Inhibitor | 4.3 | 2.0 | 0.007 |
| Spiramycin | Macrolide | Protein synthesis<br>inhibitor | 4.8 | 3.5 | 0.038 |
| cefoxitin | Cephalosporin | Cell Wall<br>Targeting | 4.0 | 3.4 | 0.005 |
| promethazine | Medication/<br>Metabolic<br>inhibitor | Repurposed<br>Medication | 4.7 | 3.3 | 0.033 |
| sodium<br>arsenate | General toxin;<br>germicide | Other | 2.1 | 2.9 | 0.032 |

a. Biolog Plate number
b. Average of WT levels with compound over WT levels in control wells. N=3
c. Average of WT levels divided by $\Delta rcsB$ mutant levels. N=3
d. Student's T-test of WT versus $\Delta rcsB$ levels.

### Enzymatic assay

Overnight cultures of wild-type bacteria were subcultured 1:100 (v/v) and grown to OD_600_= 1.65 in LB broth at 37°C in a 100 ml volume, spun down by centrifugation (7 minutes at 15,000xg), washed with an equal volume of cold PBS, and frozen at −20°C. Pellets were suspended in PBS, sonicated to lyse cells, and centrifuged (2 minutes at 15,000xg). The lysate was further clarified by passing through a 0.22 μm filter and a 10 kD size exclusion filter (Amicon, Millipore UFC901008), and washed with PBS to facilitate removal of small molecules such as D-alanine. Lysates were normalized to 23.3 mg/ml using PBS following Bradford protein analysis. Glycerol was added to 5% and the partially purified lysates were stored at −20°C or 4°C until use. The experiments were performed three times with two independent lysate preparations and results were consistent with a pilot assay using a third independent lysate.

Closely following the protocol of Garrett, et al. (67), the lysate was assessed for alanine racemase activity. The experiment was performed in three reactions, each stopped by incubation at 85°C for 10 minutes. In the first reaction the lysate (50 μL) was mixed with L-alanine to a final volume of 10 mM in PBS in a total volume of 0.1 ml, this was incubated at 37° for one hour. In this step, any alanine racemase present in the lysate could convert L-alanine to D-alanine. Negative controls for this step included no L-alanine addition and heat treatment of the lysate prior to the assay.

For the second assay the entire first reaction was added to a tube containing D-amino acid oxidase (20 units/mL, 10 μL, Sigma A5222), catalase (40 units/μL, 10 μL, Sigma C9322), FAD (1 mg/mL, 5 μL, Sigma F8384), and PBS in a total volume of 1.705 mL. This reaction proceeded for 1 hour at 37°C and was stopped by heating as above. Here, the D-amino acid oxidase converts D-alanine to pyruvate. A no D-amino acid oxidase control was included as a negative control.

The third step involved adding 1 mL of the reaction mixture with 1 mL PBS in a cuvette. NADH (10 mg/mL, 60 μL, Millipore 10128023001) was then added to all cuvettes and the mixture was allowed to incubate for 3-5 minutes. The absorbance at 340 nm was measured across a 1 cm path length in 1cm path length cuvette. Following the first A_340_ reading, lactate dehydrogenase (7200 units/mL, 10 μL, Millipore 427217) was added to all cuvettes and after 5 minutes of incubation the A_340_ was read again. The change in absorbance before and after lactate dehydrogenase addition determined the conversion of NADH to NAD^+^. Cuvettes containing known millimolar quantities of D-alanine were added to the second reaction to obtain a standard curve for comparison to reaction conditions and was linear from 0.1-100 mM of D-alanine.

To test inhibitors, the experiment was performed as above, but propionic acid was added to the first reaction at 1, 5, and 10 mM. A known alanine racemase inhibitor, β-chloro-D-alanine (0.625 mM, Sigma C4284), was used as a positive control for inhibition. Propionic acid at 10 mM was also added to the second reaction with D-alanine (10 mM) to ensure that propionic acid did not inhibit either D-amino acid oxidase or lactate dehydrogenase.

### Statistical analysis

Graphpad Prism software was used to perform ANOVA with Tukey’s post-test or Student’s T-test.

## Data Availability

The authors will provide data upon request.

## Acknowledgements

This work was supported by National Institutes of Health grants R01EY027331 (to R.S.) and CORE Grant P30 EY08098 to the Department of Ophthalmology. The Eye and Ear Foundation of Pittsburgh and from an unrestricted grant from Research to Prevent Blindness, New York, NY provided additional departmental funding.

